# Human-derived cortical neurospheroids coupled to passive, high-density and 3D MEAs: a valid platform for functional tests

**DOI:** 10.1101/2023.01.17.524439

**Authors:** L. Muzzi, D. Di Lisa, M. Falappa, S. Pepe, A. Maccione, L. Pastorino, S. Martinoia, M. Frega

## Abstract

With the advent of human-induced pluripotent stem cells (hiPSCs) differentiation protocols, different methods to create *in-vitro* human-derived neuronal networks have been proposed. Although monolayer cultures represent a valid model, adding three-dimensionality would make them more representative of an *in-vivo* environment. Thus, human-derived neurospheroids and brain-organoids are becoming increasingly used for *in-vitro* disease modeling. Achieving control over the final cell composition and investigating the exhibited electrophysiological activity is still a challenge. Thence, platforms capable of measuring and characterizing the functional aspects of these samples are needed. Here, we propose a method to rapid generate neurospheroids of human origin with control over cell composition that can be used for functional investigations. We show a characterization of the electrophysiological activity exhibited by the neurospheroids by presenting for the first-time results from the main micro-electrodes arrays (MEAs) types available on the market (passive electrode, C-MOS electrodes, 3D electrodes). Neurospheroids grown in free culture and transferred on MEA exhibited functional activity that can be chemically and electrically modulated. Our results indicates that this model holds great potential for in-depth study of signal transmission to drug screening, disease modeling and offers a reproducible and stable platform for *in-vitro* functional testing.

## Introduction

The human brain fascinates with its complexity, but the limited accessibility represents one of the major issues in the comprehension of its mechanisms. Therefore, studies of neuronal electrophysiological activity on simplified and more controlled *in-vitro* models are a fundamental step towards understanding the functioning of brain tissue. Many laboratories adopted microelectrode arrays (MEAs) technology to understand how cellular composition, connectivity, genetic and epigenetic expression correlate with the functional electrical activity expressed by *in-vitro* neuronal models(Vassallo et al., 2017, Pelkonen et al., 2021, McCready et al., 2022, Mossink et al., 2021b). MEAs applicability ranges from drug / toxicological screening (Shafer, 2019, Vassallo et al., 2017) to the characterization of various neuronal disorders (Keller and Frega, 2019, McCready et al., 2022, Mossink et al., 2021b). In addition, with the introduction of human-induced pluripotent stem cells (h-iPSCs) differentiation protocols, human-derived neuronal models could be created, potentially making the *in-vitro* approach more reliable and representative of *in-vivo* conditions. For example, this allows to generate and study neuronal cells carrying the precise genetic information of the donor in a non-invasive way easing translation of results to clinic and limiting the use of animal experimentation (Song et al., 2018, Liu et al., 2013). Although these models are widely accepted, most are based on monolayer cultures of neurons. The fact of culturing the cells only in one plane is certainly an effective simplification for many purposes, yet the *in-vitro* results are not always congruent with those *in-vivo* (Napoli and Obeid, 2016). The major difference lies in the cell morphology due to the development of flat soma and the lack of the extracellular matrix (ECM)(Duval et al., 2017).

Numerous studies aim at the realization of 3D neuronal tissues in order to obtain an *in-vitro* model that recapitulates more closely the *in-vivo* condition (Langer et al., 1995, Tang et al., 2006, Chiaradia and Lancaster, 2020). A very popular approach is to generate brain-organoids, i.e. neuronal networks derived from stem cells capable of self-organizing into a structure that resembles a human embryonic brain (Jo et al., 2016, Lancaster et al., 2013, Mariani et al., 2015, Pasca et al., 2015, Qian et al., 2016). Different approaches are adopted to favor the onset of these 3D structures, from the use of spinner vessel bioreactors (Terrasso et al., 2015, Jing and Jian-Xiong, 2011, Fair et al., 2020), matrices of round bottom wells and / or non-adherent (Jorfi et al., 2018b, Kato-Negishi et al., 2010, Boutin et al., 2018, Mukai et al., 2016), micromolding and/or microfluidics, cell aggregates on adherent substrates (Rybachuk et al., 2019, Lecomte et al., 2020, Izsak et al., 2019), hanging drop (Lee et al., 2012, Tung et al., 2011, Ganguli et al., 2021). All these approaches start from the aggregation of stem or progenitor cells, and then induce cell specialization when a 3D architecture has already been achieved. To date, several protocols are proposed that allow to guide organoids into more specialized and representative structures of certain brain regions, such as the cortex, hippocampus and midbrain (Jo et al., 2016, Mariani et al., 2015, Pasca et al., 2015, Sakaguchi et al., 2015, Yoon et al., 2019). However, these protocols generally have long timescales. Despite it has been seen how the three-dimensional organization favors the development of stem cells, the absence of vascularization in these tissues could lead to a lack of nutrients in the center of the structure, causing incomplete differentiation and / or cell death. Until now, the final result of these protocols is difficult to control in terms of cell ratio, density and makes it very challenging to create engineered and reproducible models (de Souza et al., 2018, Quadrato et al., 2017). These models are therefore currently mainly used to study the developmental trajectories of the brain during the embryonic stage (Trujillo and Muotri, 2018). Because of the variability in outcomes, the use of these models for electrophysiological investigation and generation of high throughput assays for pharmaceutical applications such as drug toxicity, drug discovery, and clinical diagnostics is still an open challenge (Jorfi et al., 2018a, Qian et al., 2019b, Tambalo and Lodato, 2020). Thus, alternative ways to generate 3D neuronal network that can be used for functional test should be investigated.

In this work, we introduce a new methodology allowing the generation of 3D neuronal networks from h-iPSCs in a relatively short time, with control over cell composition and suitable for chronical electrophysiological recordings. The proposed novel methodology is based on the combination of state-of-the art techniques to generate 3D functional neurospheroids and is based on rapid differentiation protocol for the generation of 2D population of human cortical excitatory neurons presented in Frega et al., 2017. We show the capability to grow and maintain such neurospheorids on the surface of different MEA devices commercially available for weeks. In particular we selected two planar MEAs (one based on a glass substrate providing tens of electrodes and one based on CMOS technology providing thousands of electrodes) and a 3D MEA. We demonstrate that h-iPSCs derived neurons are able to aggregate into neurospheroids with low variability in shape and without necrotic cores. Furthermore, neurospheroids exhibit spontaneous electrophysiological activity that can be modulated by receptor blockers or excited by electrical stimulation. The results indicate that this model offers a reproducible and stable platform for *in-vitro* functional testing.

## Material and Methods

### Human induced pluripotent stem cells generation and maintenance

We used a characterized rtTA/Ngn2 positive h-iPSC line generated from fibroblast of an healthy subject (30 year-old female) kindly provided in frozen vials by Frega *et al*. (Frega et al., 2017). This line was reprogrammed via episomal reprogramming (Coriell Institute for medical research, GM25256). Afterwards, rtTA/Ngn2-positive lentiviral vectors have been used to stably integrate the transgenes into the genome of the h-iPSCs. After rapid thawing, cells were plated on 6 well plate pre-coated with Matrigel (Corning) solution (1:15 in cold Dulbecco’s Modified Eagle Medium (DMEM, Gibco, Thermofisher scientific inc.)) and cultured in Essential 8 Flex (E8F) Medium (Thermo Fisher scientific) supplemented with Essential 8 Flex supplement and 1% pen/strep (Thermofisher Scientific). The medium was also supplemented with 50 μg/ml G418 (Sigma-Aldrich) and 0.5 μg/ml puromycin (Sigma-Aldrich), selective antibiotic to preserve the purity of the cell line. Medium was refreshed every 2–3 days and cells were passaged twice per week using an enzyme-free reagent (ReLeSR, Stem Cell Technologies).

### Neuronal differentiation and neurospheroids generation

h-iPSCs were directly derived into excitatory cortical Layer 2/3 neurons by overexpressing the neuronal determinant Neurogenin 2 (*Ngn2*) upon doxycycline treatment as described previously (Frega et al., 2017). In particular, h-iPSCs were detached from a well after reaching confluence (6 × 10^6^ cells) using ReleSR and a single-cell solution was obtained by collecting the colonies in 2 ml of “E8F+dox” medium (i.e. E8F medium supplemented with Essential 8 Flex supplement, 1% pen/strep, 50 μg/ml G418, 0.5 μg/ml puromycin and 4 µg/m doxycycline (Sigma-Aldrich)). Consequently, we added 1 ml of single-cell solution to Matrigel pre-coated wells in which 1 ml of “E8F+dox” medium was previously plated and we stored the plate into the incubator (Figure 1a, Day 0). The next day (Figure 1a, Day 1) we replaced the “E8F+dox” medium with DMEM/F12 (Gibco) supplemented with MEM non-essential amino acid solution 100x (Sigma Aldrich), 100x N2-supplement (Invitrogen), 100x pens/strep, 10 ng/ml human-NT-3 (BioConnect), 10 ng/ml human-BDNF (BioConnect) and 1% pen/strep. At Day 3 cells (i.e. “early-stage neurons”) have reached confluence and are ready to be detached and used in co-culture with astrocytes. In particular, early-stage neurons were detached using Accutase (5 minutes at 37°C, StemPro) and transferred into a 15 ml flacon tube containing 8 ml of Neurobasal medium (Gibco) supplemented with B-27 Supplement 50x (Gibco), GlutaMax 100x (Invitrogen), 5 μg/ml Doxycycline, 10 ng/ml h-BDNF 10 ng/ml h-NT-3, 100x pen/strep. After centrifugation (1200 rpm for 5 min), the supernatant was removed and the early-stage neurons were suspended in 2 ml of medium. Early-stage neurons are now ready to be counted and used in co-culture for the neurospheroids generation (Figure 1b).

**Figure 1.**
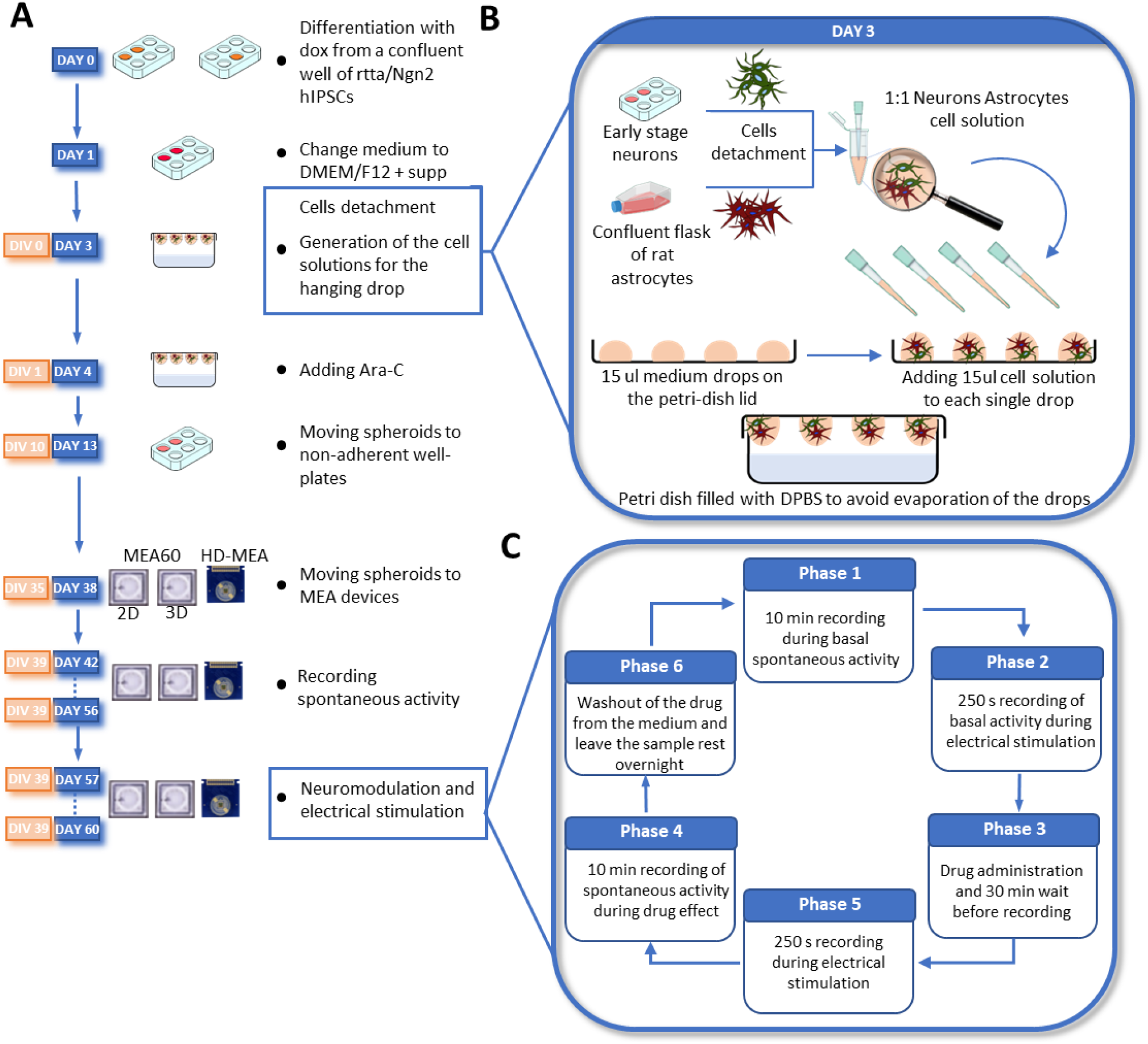
Overview of the differentiation steps and experimental design. **a)** Timeline of the development of the neurospherois from the differentiation process to the transfer and growth onto the MEAs. **b)** Focus on day 3/DIV 0 main steps, collection of cells and hanging drop technique. Early-stage neurons and astrocytes are harvested the same day in two separate tubes. Consequently, cells are counted and mixed in a 1:1 ratio in a 1 ml vial at a final concentration of 5’330’000 cell/ml. Therefore, we plated 15 µl of the cell suspension on the 15 µl medium drops in the lid of a petri-dish. Finally, we gently placed back the lid on the petri-dish bottom part half-filled with DPBS and then we stored everything in the incubator. **c)** Experimental protocol for chemical neuromodulation and electrical stimulation.

Neurospheroids were composed by 80’0000 early-stage neurons and astrocytes in 1:1 ratio. Astrocytes were obtained from brain cortices of day 18 embryos as described in Aprile *et al*. (Aprile et al., 2019) and are needed to support neuronal maturation. We choose to exploit the hanging drop method to generate spherical aggregates of cells and we used a 5 cm petri dish as ‘moisture chamber’ for the neurospheroids. We half-filled the petri dish with DPBS and we used the inner part of the lid as sustain for the drops of medium used as scaffold-free culture. In detail, we inverted the lid and we placed eight 15 µl drops of neurobasal medium into the inner part of the lid. Thereafter, we added 15 µl of the mixed-cells solution (i.e. solution of 5’330’000 cell/ml composed by 500’000 early neurons and 500’000 astrocytes) to each medium drops previously placed. Consequently, we placed the lid back on the petri dish with a gentle but rapid movement to avoid sliding of the drops. We then stored the dish into the incubator at 37°C and 5,5 C0_2_. We consider this day as Day in Vitro (DIV 0) of the neurospheroids. On DIV 1 we added Ara-C (Sigma-Aldrich) to each drop to a final concentration of 2 µM (Figure 1a, day 4). During the first week *in vitro*, we added medium if evaporation occurred. From DIV 7 on, we replaced 10 µl of medium for each drop, paying attention to avoid damaging the forming neurospheroids. At DIV 10, neurospheroids were moved into 24 well-plates pre-coated with 1% w/v Alginate solution to avoid cell attachment and preserve the free-culture. From this moment on, medium was supplement with 2,5% of FBS and 30% of the medium was refreshed three times a week. At DIV 35, neurospheroids were moved into MEAs.

### MEA devices

To evaluate the functional activity of the neurospheroids, we used MEA60 (Multi Channel System, MCS, GmbH), 3D MEA60 (Multi Channel System, MCS, GmbH) and Accura High Density MEA (HD-MEA) from 3Brain AG, Switzerland. The MEA60 are glass devices where 60 titanium nitride electrodes are embedded in the center of the culture well. They are arranged in a 8×8 grid without electrodes at the corner, spaced 200 µm among them with 30 µm diameter, generating an active area of 1,6 mm x 1,6 mm. The 3D MEA60 have the same spatial organization of the electrodes, with the difference that electrodes are 250 µm spaced among them and are pyramidal with 100 µm height and 12 µm diameter tip. The HD-MEA (Accura model, 3Brain AG) devices present an active area of 3,8 mm x 3,8 mm in which 4096 CMOS electrodes are integrated. Each microelectrode has a dimension of 21 µm x 21 µm with a 60 µm pitch, arranged in a 64×64 grid.

Prior to the use, each device was cleaned and sterilized. In particular, MEA60 and 3D MEA60 were sterilized in the hoven at 120°C for 2h. The HD-MEAs were cleaned with DPBS and the culture chamber was filled with 70% EtOH for 20 min for sterilization. After ethanol, the chambers were washed tree times with DPBS. Then, each chamber was filled with 2ml of DPBS and the devices were stored in the incubator for 2 nights, to improve hydrophilicity of the substrate (conditioning phase). After this phase, all the devices were coated with a bi-layer composed of poly-L-ornithine (PLO, Sigma-Aldrich) and human-laminin (BioConnect). More specifically, we deposited a 100 µl drop on the active area from a fresh solution of 50 µg/ml PLO in DBPS and we incubated the devices at 4°C overnight. The day after, we removed the drop containing PLO, we washed twice the chambers with DPBS and we deposited an 80 µl drop of 20 µg/ml human laminin onto the active area. Devices were then left overnight at 4°C, to be used the subsequent day.

### Data acquisition and analysis

We recorded 10 minutes of spontaneous activity at DIV 42-49-56 using the 2100 System (MEA 2100-System,MCS) for the MEA60 and 3D MEA60 and the BioCam Duplex (3Brain AG) for the HD-MEAs. Data were sampled at 20 kHz. Incubator-like conditions were maintained during recording by keeping the culture at 37°C and 5.5% CO_2_ in sterile conditions. Cultures were also recorded at DIV 57 and 60 to perform electrical ad chemical stimulation (Figure 1c).

Data recorded from MEA60 (and 3D MEA60) were analyzed using custom Matlab scripts, prior conversion to *hdf5* format using MultiChannel DataManager (MCS). Regarding the HD-MEA, BrainWave software (v.4.4, 3Brain AG) was used to high-pass filter the signal at 200 Hz using a 2^nd^ order Butterworth filter and for the PTSD. Consequently, we used custom Matlab scripts to read results from Precise-Timing Spike Detection (PTSD) and the analysis proceed with Matlab script.

#### Spike detection

The PTSD algorithm was used to detect the spikes (Maccione et al., 2009) Standard deviation factor (TH_sd_) of 6 and 8 for passive MEAs and HD-MEA respectively, peak-lifetime period of 1ms, refractory period of 1ms and spike (spk) assigned to the higher peak were used as detection settings.

#### Mean firing rate

The Mean Firing Rate (MFR) was evaluated the for each electrode as the ratio between all the detected spikes and the recording time. We considered into the analysis only the ‘active’ electrodes exhibiting a MFR>0,1 spike/s.

#### Burst detection

we considered a burst as a series of 3 consecutive spikes firing no more than 50 ms apart from each other. The Mean Burst Rate (MBR) was evaluated as the sum of all the detected burst occurred in an active channel divided the recording time, and we considered bursting electrodes those having a MBR>0,2 burst/min. Mean Burst Duration (MBD) was evaluated by averaging all the burst duration detected in the whole culture. Therefore, we evaluated the percentage of random spikes (PRS) as the ratio between total number of non-burst spikes and total number of spikes. Similarly, we evaluated the percentage of bursting channels (PBC) as the ratio between bursting electrodes and active electrodes.

#### Raster plots

We generated rasterplots to show an overview of the total activity being recorded from each electrodes of the MEA. We represent the activity (i.e. dots indicating a spike detected) recorded from each included recording electrode (i.e. single line on the y-axis) during time (i.e. x-axis).

#### Network burst detection

Synchronous events occurring within the network, defined as *Network Bursts* (NBs) were detected as previously reported in (Muzzi et al 2021).

#### Post-Stimulus Time Histogram

To evaluate the electrical response, we computed the Post-Stimulus Time Histogram (PSTH) by counting all the detected spikes occurring after each stimulation in 10 ms bin. We then quantified the fast response by counting all the evocated spikes occurring in the first 100 ms, the late response as the spikes occurred between 100 ms and 1000 ms and the peak latency as the time of the maximum activation. To normalize the PSTH, we obtained the Pre-stimulus time histograms by counting the detected spikes occurred 200 ms before each stimulation in 10 ms bins. We choose the mean value across bins in the Pre-stimulus time Histogram to obtain a normalization factor for each PSTH. Each normalized PSTH was then averaged across the 50-stimulation delivered in that culture and average across neurospheroids cultured on same MEAs.

#### Cross-Correlation

To evaluate possible functional connections between electrodes, we implemented a Cross-Correlation (CC) script in Matlab using a common normalization factor (Eytan et al., 2004). The cross correlation expresses the probability Cxy(τ) of observing a spike in one electrode y at time (t + τ), given that there was a spike in a reference electrode x at time t. We calculated the connectivity matrices by analyzing the correlograms obtained in a time window of 300 ms centered at time 0 and using a 0.2 ms bin. In order to eliminate the contribution of spurious connections we have applied a threshold to filter the connections based on their weight. The threshold was calculated as the average of the weights of the total connections plus 2 times the value of the standard deviation. Consequently, for each matrix we calculated the total number of links, their average weight and the average connection time τ (i.e. time lag). To reduce computational time we applied the CC algorithm to 144 electrodes arranged in a 12 × 12 square grid placed under the neurospheroid.

#### Single cell signal propagation

Taking advantage of the high spatial resolution electrode array of the HD-MEA, we performed an analysis on the velocity of propagation of spikes along neuronal axons. To obtain this, we averaged raw data traces over multiple spike occurrences of the same neuron. After spike sorting based on PCA and k-means clustering, individual units (i.e., neurons) with an average peak amplitude of at least 300 µV were taken into account. For each considered unit, we computed the spike timestamp and we extracted the raw traces of the entire electrode array in a time window of 10 ms around each spike. We then averaged all the extracted raw data traces to obtain the averaged signal propagation and we calculated the traveled distance of each propagation. To do this we identified the electrodes involved and we calculated the total distance as the sum of the distances between consecutive electrodes involved in the propagation. Consequently, we calculated the average speed by taking the time difference between the first and last spike of the propagation divided by the total distance calculated as explained above.

### Electrical stimulation and neuromodulation

At DIV 57 functional tests on the neurospheroids were performed (Figure 1c). We evaluated the effect of 3 neuromodulators, both in terms of spontaneous activity and in response to an electrical stimulation. The protocol we used includes 10 minutes of recording in spontaneous conditions, followed by a current stimulation protocol. Consequently, the cultures were treated with one of the neuromodulators and placed in the incubator for 30 minutes. After this time, we recorded 10 minutes of spontaneous activity in presence of drug and then we applied the stimulation protocol again. Finally, we removed the neuromodulator from the sample by 3 consecutive washing steps with a previously conditioned medium and left the cultures to rest for at least 12 hours before testing the new drug. The neuromodulators used are 6-cyano-7-nitroquinoxaline-2,3-dione (CNQX, Sigma-Aldrich) 50 µM, D-(–)-2-Amino-5-phosphonopentanoic Acid (D-APV, Sigma-Aldrich) 80 µM, Kainic Acid (KA, Sigma-Aldrich) 5 µM. Stimulation protocol consist of injecting 50 bipolar current pulses of 60 µA peak-to-peak amplitude, half-width of 100 µs, at 0,1 Hz from a pair of electrodes selected under the neurospheroids.

### Immunofluorescence

After the electrophysiological recordings to characterize the neurospheroids development and neuronal response to electrical stimulation and neuromodulators, samples were fixed directly on the HD-MEA at DIV 60, and fluorescence images were acquired. For the fixation, the samples were exposed to 4% paraformaldehyde solution (PFA, Sigma-Aldrich) for 20 min at room temperature and then washed 3 times in Phosphate-buffered saline (PBS, Sigma-Aldrich) solution. Samples were permeabilized with 0,2% triton X-100 (Thermo Fisher Scientific) for 15 min. To block non-specific binding antibodies, samples were exposed to Blocking Buffer Solution (BBS, composed with 0.5% fetal bovine serum (Sigma-Aldrich), 0.3% bovine serum albumin (Sigma-Aldrich) in PBS) for 45 minutes at room temperature. We used GFAP (diluted 1:500, Sigma Aldrich) and MAP-2 (diluted 1:500, Chemicon Millipore) as primary antibodies to mark glial and neuronal cells, respectively, and Dapi (diluted 1:10000, Sigma) to label nuclei. We used Alexa Fluor 488 (diluted 1:700, Thermo Fisher Scientific) and Alexa Fluor 546 (diluted 1:1000, Invitrogen) Goat anti mouse or Goat anti rabbit as secondary antibodies.

### Statistical analysis

In this work we show data obtained from n=2 MEA60, n=2 3D MEA, n=2 3D HD-MEA. Statistical analysis was performed on data obtained from all active electrodes. After evaluation of normality test carried out with GraphPad Prism we performed non parametrical Wilcoxon signed-rank test. Differences were considered significant when p<0.05.

## Results

### Human-iPSCs derived neurons aggregate into neurospheroids

First, we monitored the formation of the neurospheroids during time and we observed that h-iPSCs derived neurons started aggregating after about 10 days of culturing in the petri dish. When moved into free-cultures, the aggregates showed a spherical shape with a mean diameter of 412 ± 16 µm and 429 ± 20 µm at DIV 40 and DIV 62, respectively (Figure 2a,b). Then, we investigated the development of the neurospheroids on MEAs. Neurospheroids adhered to all MEAs and we observed that the height of the structure was lower than the mean diameter measured from the samples kept in free culture (Figure 2c). In particular, we found that the highest point of the neurospheroids growing on a HD-MEA is at 180 µm from the electrode plane and that the diameter was ∼550 µm (Figure 2c,d). Furthermore, we observed that both neurons and astrocytes were present within the neurospheroids and that cells were viable in the inner part since nuclei were round and regular through the entire height of the structure (Figure 2c,d and Supplementary videos 1).

**Figure 2.**
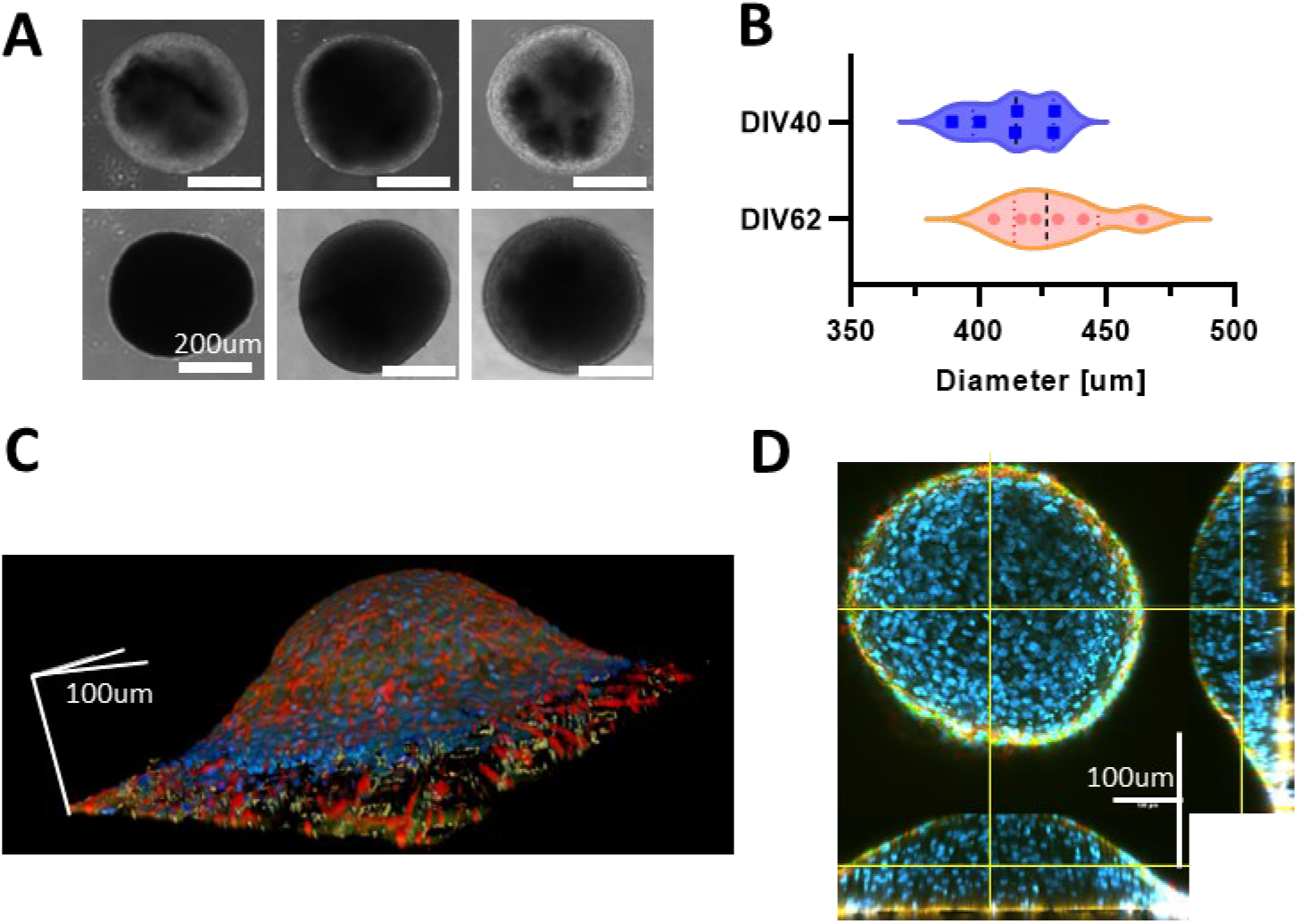
**a)** Images of different neurospheroids in free-culture during development. First row shows picture obtained at DIV 40 and second row at DIV 62. Scale bar 200 µm. **b)** Evaluation of the mean diameter of N=6 neurospheroid at DIV 40 and DIV 62 in free culture. **c)** Reconstruction of the 3D volume of a neurospheroid plated on a HD-MEA at DIV 35 and fixed at DIV 62. Images were acquired with a confocal microscope using a z-stack of Scale bars represent 100 µm in each direction. **d)** Orthogonal view of the sample shown in b. Yellow lines represent the observation plane. In Blue: DAPI, Green: MAP2, red: GFAP. Scale bar represent 100 µm in each direction

### Neurospheroids showed mature electrophysiological activity on MEAs

Neurospheorids were plated on the three different MEA types (Figure3a-c) at DIV35 to compare the functional activity recorded by the three devices. We observed that neurospheroids exhibited mature electrophysiological activity that could be detected by all devices at DIV 42. By comparing the raw data of a single electrode under the neurospheroids, we observed different levels of noise and spike amplitude. In particular, HD-MEA detects significantly larger waveforms as compared to the lower resolution devices. This fact combined with the higher resolution of the HD-MEA, allows to clearly identify the onset of network events (i.e. darker vertical bands in raster plots, Figure 3 g-i).

**Figure 3.**
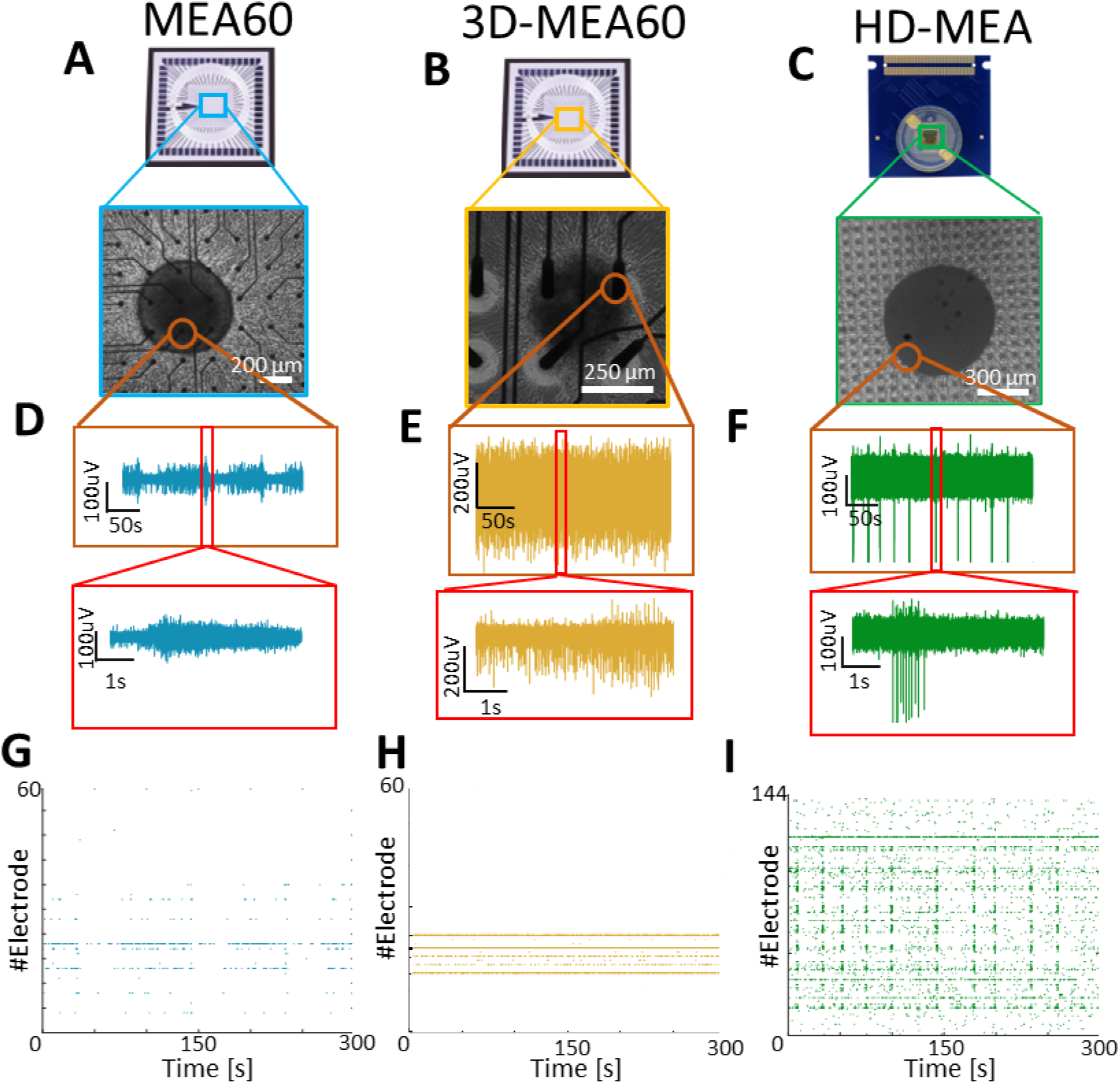
Qualitative comparison between activity recorded by MEA60, 3D-MEA60 and HD-MEA. **a-c)** Picture showing a neurospheroid at DIV 49 on **a)** MEA60 (scale bar 200 µm), **b)** 3D-MEA60 (scale bar 200 µm), and **c)** HD-MEA (scale bar 300 µm). **d-f)** Exemplificative raw data trace recorded by one electrode placed below the neurospheroid growing on **d)** MEA60, **e)** 3D-MEA60, and **f)** HD-MEA. 5 minutes and 5 second close-up of the activity are shown in the orange and red box respectively. **g-i)** Raster plot showing 5 minutes of activity exhibited by a neurospheroid recorded by **g)** MEA60, **h)** 3D-MEA60, and **i)** HD-MEA. Each dot represents a detected spikes at the given electrode index (y axis).

We then followed the development of the neurospheroids by recording the spontaneous activity at different time point (DIV 42—49 – 56). We found that the number of active electrodes and most of the firing metrics tends to increase over time in neurospheroids plated on HD-MEAs while they remain almost stable in the MEA60s (Figure 4a-d). In all cases, we observed that activity evolved over time, moving from mainly random spiking to bursting. Indeed, percentage of random spikes (PRS) decreased over time while the percentage of active bursting channels increased (Figure 4g-h). With all devices we were able to detect phenomena of synchronous network activity, although with the MEA60 and 3D-MEA60 they only emerged later in development (i.e. from DIV 49 on) and with a significantly lower frequency as compared to HD-MEA (Figure 4e,i-k). Even if the duration of the detected NBs was variable between the different devices (Figure 4f), the average shapes of the NBs were similar for the three devices, but are very scattered in the case of the MEA60 and 3D-MEA60 (Figure 4l).

**Figure 4.**
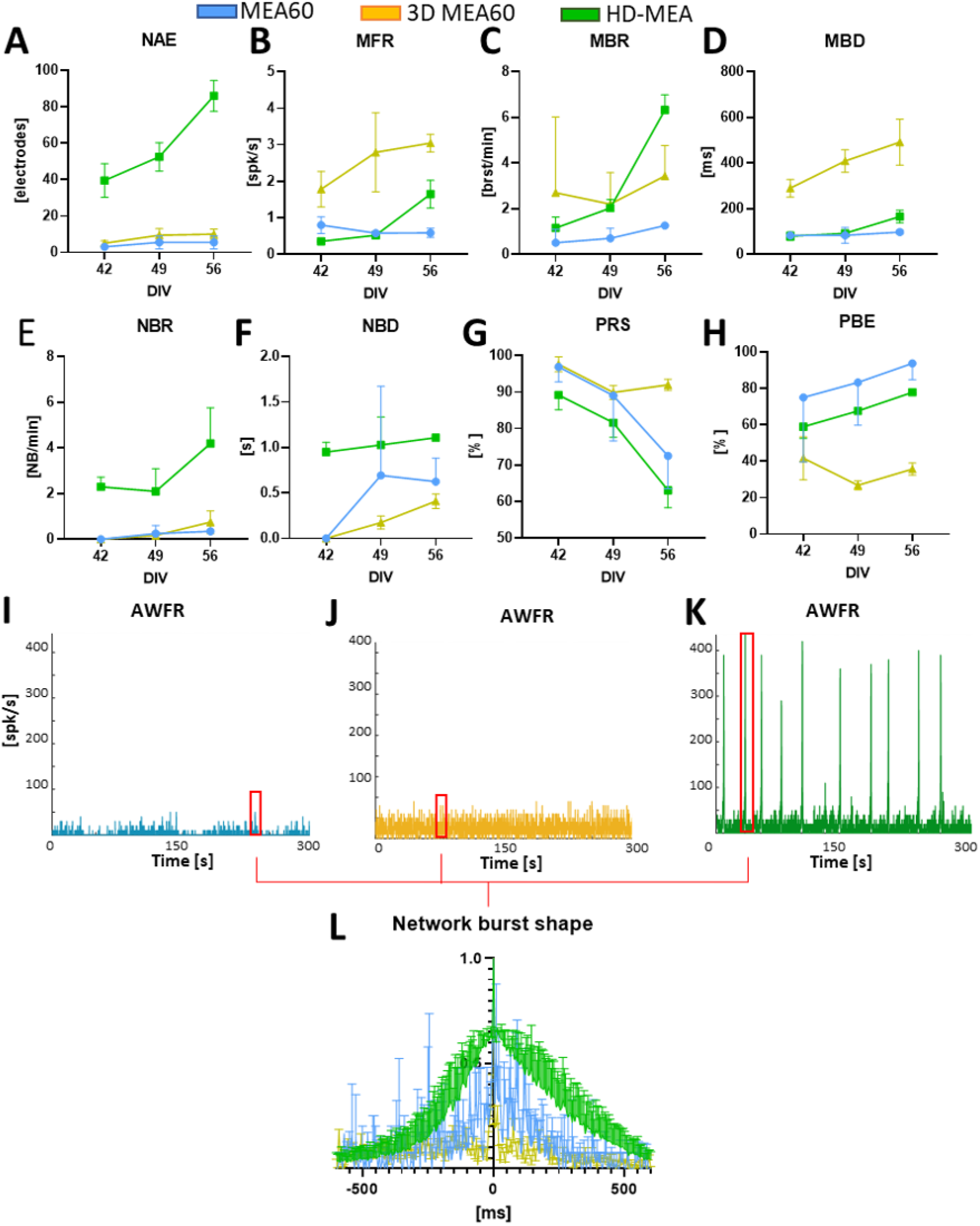
Analysis of the spontaneous activity of n=2 samples on MEA60 (data showed in blue), n=2 HD-MEA (data showed in green), n=2 3D-MEA (data showed in yellow). **a-h)** Graphs showing the **a)** number of active electrodes (NAE), **b)** Mean firing rate, **c)**, Mean bursting rate (MBR), **d)** Mean burst duration (MBD), **e)** Network burst rate (NBR), **f)** Network burst duration (NBD), **g)** Percentual random spikes (PRS), **h)** Percentual bursting channels (PBC). Data are shown as mean and standard deviation of the mean. **i-k)** Graphs showing the Array Wide Firing Rate in neurospheroids growing on **i)** MEA60, **j)** 3D-MEA60 and **k)** HD-MEA. It quantifies the level of activity as averaged firing rate of the all network evaluated in 100 ms bins. Network Burst (NB) are highlighted by red rectangles. **l)** Mean NB shape aligned and normalized by the higher peak.

### Neuromodulation affects spontaneous activity and electrical induced activity

Once neurospheroids reached a steady-state regime (i.e. presence of robust Network Bursts) at DIV 57, we subjected them to electrical stimulation conjugated with chemical modulation protocols in order to evaluate the effects of three neuromodulators on functional activity (Figure 5 and Supplementary Figure 1). The inhibition of the AMPA receptors (CNQX 50 µM) led to a significant reduction of the mean firing rate and bursting rate (Figure 5c,d). In addition, the synchronous network activity was suppressed (Figure 5f,g) causing an increase in random activity and a significant decrease in active and bursting electrodes (Figure 5a,h,i). As regards the effect on electrical stimulation, the response to stimuli in the presence of CNQX is slower as compared to basal condition (Figure 5j, peak latency) with a significant reduction of evoked potentials in the first 100 ms following the application of the stimulus. The suppression of NMDA receptors (APV 80 µM) did not affect the number of active electrodes and did not significantly reduce the mean firing rate and mean bursting rate (Figure 5 c,d). The synchronous activity also remained almost unchanged while the random activity generally decreased (Figure 5 f,h). By evaluating the change in response to electrical stimulation, we found a significant delay in reaching the maximum response peak and a decrease in spikes evoked in the first 100 ms (figure 5k). Finally, Kainic acid (5 µM) had a significant effect on the synchronous network activity, increasing its rate while maintaining a similar average duration of events (Figure 5 f,g). There was also a substantial decrease in active electrodes. Evaluating the response to the stimulus, the results showed an effect similar to that caused by CNQX, with an anticipation of the peak and a reduction in rapid evoked potentials, but it did not show significant differences regarding the late response phase (Figure 5k).

**Figure 5.**
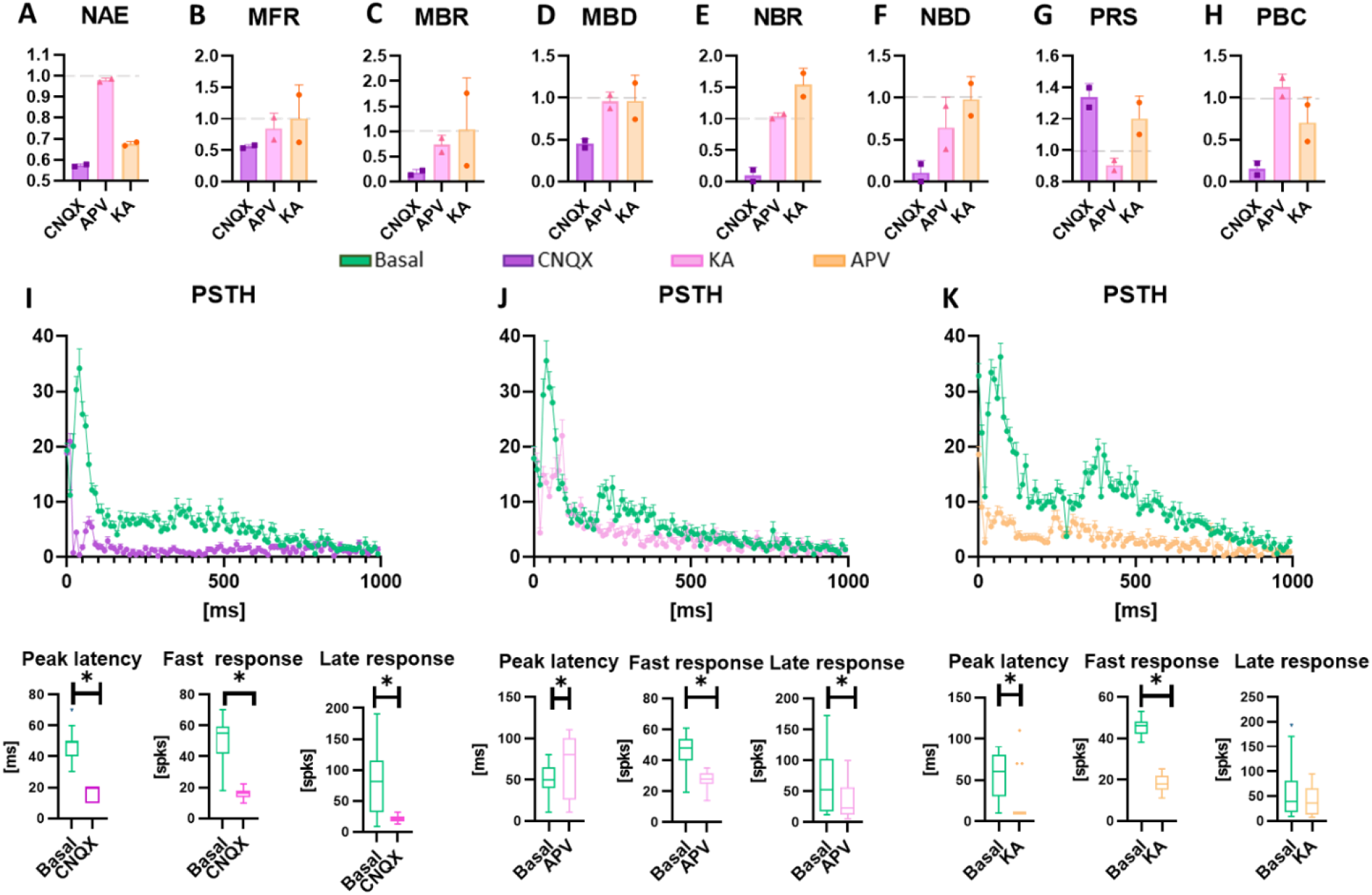
Evaluation of the electrical and chemical modulation (50 µM CNQX, 80 µM APV, and 5 µM KA). Data are obtained from experiments on HD-MEA (n = 2) and are normalized to the value detected in spontaneous conditions in the absence of drugs. Data obtained with MEA60 are reported in Supplementary Figure 1. **a-h)** Graphs showing the **a)** Number of active electrodes (NAE), **b)** Mean firing rate (MFR), **c)** Mean bursting Rate (MBR), **d)** Mean burst duration (MBD), **e)** Network Burst rate (NBR), **f)** Network Burst duration (NBD), **g)** Percentual random spikes (PRS), **h)** Percentual bursting channels (PBC) in neurospheroids treated with CNQX, APC and KA, represented int purple, pink and orange, respectively. Data are represented as mean and standard deviation of the mean. **i-k)** Graphs showing the effect of the modulation on the response to the electrical stimulus induced by **i)** CNQX, **j)** APV, **k)** KA. In each panel the Post-Stimulus Time Histogram (PSTH) showing the response of neurospheroids to stimulation is shown (not treated, treated with CNQX, APV and KA are represented in green, purple, pink and orange, respectively). The peak latency represent the mean latency to reach the maximum peak in the PSTHs, the fast response box plot represents the amount of evocated spikes in the first 100 ms of the PSTH while the late response box represent the amount of evocated spikes between 100 ms and 1000 ms. Asterisk indicate p<0.05.

### Neuromodulation affects connectivity and signal propagation

To evaluate how neuromodulation and electrical stimulation affected the connectivity of the neurospheroids we performed cross-correlation analysis. We evaluate the CC of the neurospheroids under several condition. First, results indicate that inhibition of AMPA receptors with CNQX led to a decrease in the number of links and increase in the mean weight (Figure 6 b,c), while electrical stimulation led to an increase of the number of links and a slightly decrease in the mean weight (Figure 6 e,f). Comparison between electrical stimulation in spontaneous condition and CNQX stimulation led to a decrease in total number of links and increase in their mean weight (Figure 6 h,i). The blockage of NMDA receptors with APV caused a less significative change of connectivity in the absence of stimulation. The application of electrical stimulation led to a decrease of the number of links, an increase of their mean weight and a slightly reduction of the mean lag when compared to electrical stimulation in basal condition. Electrical stimulation under APV effect resulted in decreasing the number of links, increasing their mean weight and slightly decreasing their mean lag compared to electrical stimulation in basal condition (Figure 6 h-j). Finally, electrical stimulation after KA administration led to a decrease in the number of links and a slight increase of their mean weight.

**Figure 6.**
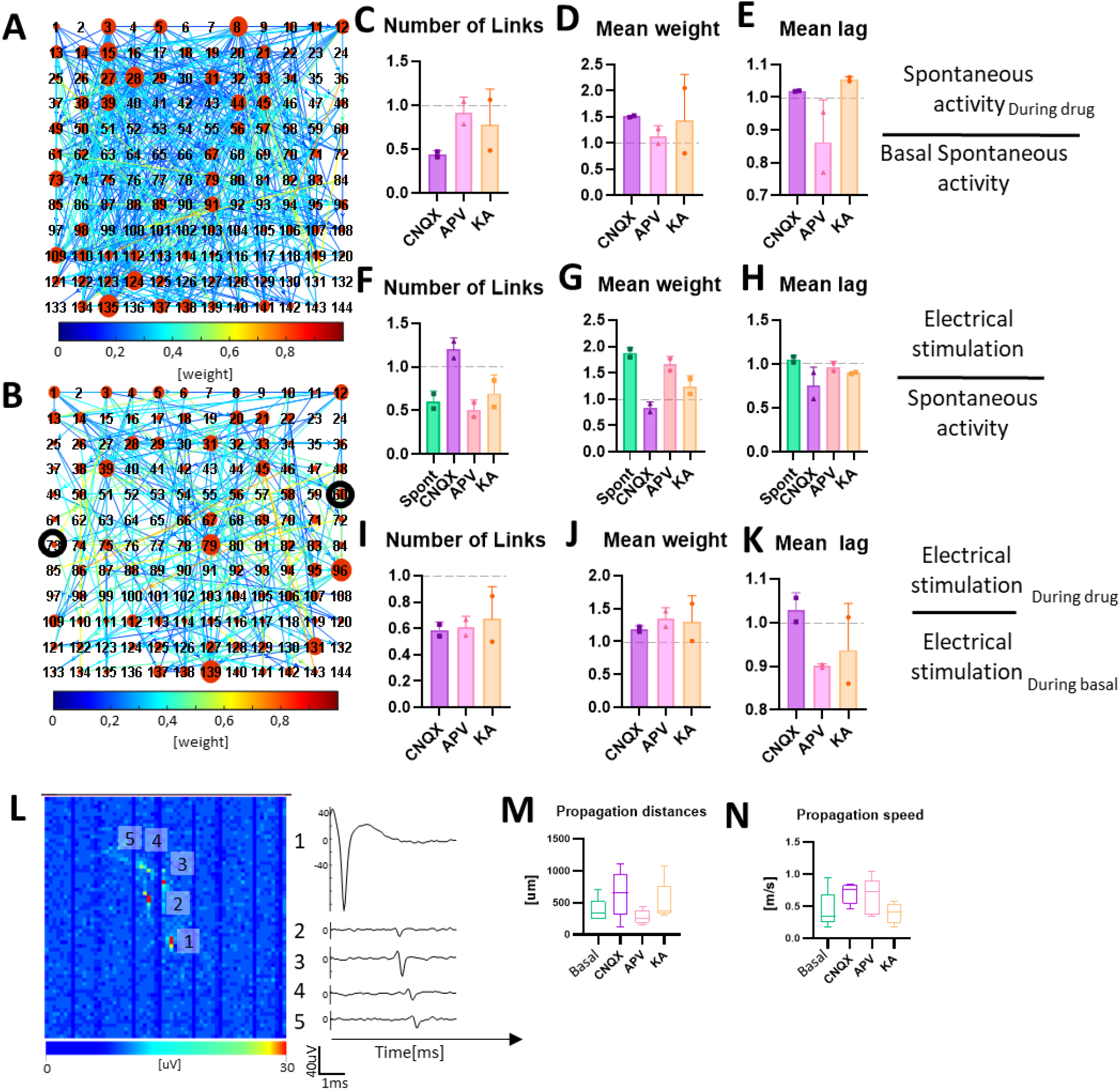
Neuronal connectivity in neurospheroids. **a)** Connectivity matrix obtained from a neurospheroid grown on HD-MEA at DIV 57 during spontaneous basal activity and **b)** during electrical stimulation. Each red circle represents an electrode, the larger the size of the red circle the greater the number of links. Colormap represent the weight of each link. Darker circle indicates electrodes used for delivering stimulation. We restricted the connectivity analysis to 144 electrodes arranged in a 12 × 12 square grid placed under the neurospheroid. 12 **c-e)** Graphs showing the effect of neuromodulators on connectivity, in particular: **c)** number of links, **d)** mean weight and **e)** mean lag evaluated on 10 minutes of activity under the effects of the drugs, normalized on 10 minutes of spontaneous electrical activity without drug. **f-h)** Graphs showing the effect of electrical stimulation during neuromodulation on connectivity, in particular: **f)** number of links, **g)** mean weight and **h)** mean lag evaluated on 10 minutes of recording during electrical stimulation under the effects of the drugs (or no drug for electrical stimulation, labelled as basal), normalized with data obtained from the recording of spontaneous activity in the same conditions. **i-k)** Graphs shows the overall effect on connectivity of the electrical stimulation during neuromodulation, in particular: **i)** number of links, **j)** mean weight and **k)** mean lag evaluated during the electrical stimulation under the effect of the neuromodulator, normalized by the data obtained from recording during electrical stimulation without drugs. Bars represent mean value, point represent single experiments and errors are expressed using standard deviation. **l**) On the left, Activity map of the HD-MEA showing the averaged axonal propagation of a single neuron. The image is composed of 64 by 64 pixels, where each pixel represents the voltage amplitude of an electrode. Scale bar represent maximum variation of voltage in a 1 ms sliding window. On the right, voltage traces of five electrodes recording the signal propagation.. Numbers identifies the electrodes selected on the activity map. **m-n)** Graphs showing **m)** mean propagation distance and **n)** mean propagation speed of ten neurons in basal and chemically stimulated conditions.

Thanks to the high spatial resolution offered by the HD-MEA devices, it was possible to highlight, trace and estimate the speed of action potentials in the various experimental phases (Figure 6l-n). Under spontaneous conditions we found propagations covering mean lengths of 375 ± 194 µm with mean speeds of 0.44 ± 0.30 m/s (Figure 6 m,n). In the presence of CNQX, we calculated higher speeds with lower variability (0.7 ± 0.15 m/s) but over wider mean distances 636 ± 362 µm. In the presence of APV we found shorter propagations (278 ± 112 µm), with an average velocity more similar to that estimated with CNQX but more variable (0.65 ± 0.29 m/s). Finally, with KA we found average propagation lengths of 510 ± 300 µm with comparable and less varied velocity than in the normal condition (0.39 ± 0.16 m/s).

## Discussion

In this article we presented a protocol to generate functional cortical excitatory human neurospheroids suitable for *in-vitro* functional investigation through MEA devices. We used a protocol based on Ngn2 induction as it allows the generation of a homogeneous population of cortical excitatory neurons that are mature and functional in a short time (i.e., 5 weeks, Frega et al., 2019). Furthermore, we improved the protocol, to obtain an increased control over the plating densities (Muzzi et al., 2021b). Notably this methodology can be used in combination with other rapid differentiation protocols to generate other neuronal types (Mossink et al., 2021a, Yang et al., 2017) offering a broad spectrum of designability for 2D and 3D tissues.

The use of the hanging-drop technique made it possible to generate the neurospheroids without the use of bioreactors or scaffold. We have noticed that the cells needed 7 to 10 days to form a spheroidal structure inside the drop. After 10 days, all the drops in the petri dishes contained solid and regular structures so we decided to move them to non-adherent multi-well plates. The main reason is due to the difficult maintenance of the neurospheroids in the hanging-drop condition. In particular, cells have small volume of medium available, they are easily subjected to evaporation and mechanical stresses could damage the viability of the 3D network. Once the structures were transferred to the plates, their maintenance proved to be much easier, however they still need to be stirred from time to time to avoid to become oval. Although we did not find significant size differences between the generated samples, we did notice a slight increase in size over time (see Figure 2b). This might be due to a proliferating astrocytic part, which recovered after the initial phases of aggregation of the neurospheroids where we treated it with Ara-C. The shapes of the nuclei in the core of the neurospheroids are round and regular (from the z-stack) indicating that cells are healthy. This is an essential point in neurospheroids generation in which the core represents the most sensitive part because it is subjected to necrosis due to the lack of nutrient exchange (Qian et al., 2019a). We also noticed that the samples on the MEA devices adapted their dimensions and preferred a radial growth on the substrate rather than preserving the spherical shape. In fact, after 30 days in culture we found a maximum height of the neurospheroids of 180 µm with a diameter of about 550 µm, thus suggesting a flattening of the structure on the electrode plane, also confirmed by the increased number of active electrodes in the MEAs out of the spheroid area.

To verify the functionality of the neurospheroids, we monitored the development of spontaneous electrophysiological activity for 5 weeks (from the moment of adhesion to the device) on 3 different MEA devices. This allows us to investigate whether coupling these structures with MEA devices could constitute a valid 3D *in-vitro* model useful for functional characterizing. We used two types of MEA with passive electrodes, one with planar (MEA60) and one with 3D electrodes penetrating the structure and recording signals internally (3D-MEAs). In addition, to achieve a high-density spatial resolution for the recording of the electrophysiological activity and a higher signal to noise ratio, we used an advanced device that exploit the CMOS technology (HD-MEA). We moved the culture to the devices at DIV 35 because at this stage of development a steady-state and stable network dynamic had been observed in 2D and 3D (Frega et al., 2019b, Muzzi et al., 2021a). During the first few days after adhesion, we were unable to record activity. This was probably due to the fact that cells must adhere to the electrode plane to develop synapses and functional connections before a signal can be picked up. After a week, we found active electrodes in all the devices and already a quite mature network dynamic, presenting single channel activity and burst. During time, the active bursting channels increase and PRS decrease, showing trends in line with the results of others 3D cultures in the literature (Tedesco et al., 2018, Frega et al., 2014, Izsak et al., 2019). During the recording weeks, several electrodes became active even far away from the neurospheroids. Many cells moved from the neurospheroids to the electrode plane creating a peripheral 2D network from which activity can be detected (see Figure 2). We observed differences in levels of firing rate recorded from different MEAs. This derives from the different nature and density of the electrodes. In particular, the raw signal recorded by MEA60s appears noisier as compared to HD-MEAs making difficult to detect close spikes in moments of high synchronous activity such as NBs. Consequently, lower electrode density combined with a lower Signal / Noise ratio led to a different quantification of the MBR, MBD and NBR in the passive MEAs as compared to HD-MEAs. Similar considerations apply to 3D-MEAs, with the difference that the active electrodes are located inside the neurospheroids and can therefore record activity by cells in all directions. The presence of electrophysiological signal and the fact that we are able to detect NB is a further confirmation of the potential of these cultures to be used for functional tests. Once we verified that the neurospheroids were active and mature in electrophysiological terms, both internally and externally, we evaluated their response to electrical stimuli in conjunction with neuromodulators. Although we have performed these experiments on all the samples in each type of MEA, we show in the main text only the data obtained from the HD-MEAs. Indeed, HD-MEAs resulted the most suitable device to record electrophysiological activity thanks to the higher spatial resolution and better signal to noise ratio acquisition allowing a more accurate analysis. Interestingly, neurospheroids on 3D-MEAs detached after the application of the first stimulation protocol. This might be caused by variations in the electrode impedance due to the production process and the fine tip, which can alter stimulation intensities, especially with currents (Systems, 2021). It is therefore possible that the actual stimulation current resulted higher than the set value, causing irremediable damage to the neurospheroids.

Since the neurospheroids are composed only of excitatory neurons we have used neuromodulators that interact with glutamatergic receptors. CNQX and APV respectively inhibit AMPA and NMDA receptors which we know are the main mediators of the network activity of these culture (Frega et al., 2019).The administration of CNQX led to a radical change in the network dynamics of the neurospheroids. The spontaneous AMPA-inhibited activity showed a significant decrease in network properties compared to baseline, almost completely suppressing single-channel and network bursting activity. The inhibition of NMDA receptors alone did not affect significantly the spontaneous activity of the neurospheroids. The effect of the modulator is observed in a decrease of about 40% in the percentage of random spikes and a reduction in the average length of the NBs. This reflects the results of previous studies (Durens et al., 2020, Trujillo et al., 2019) showing that NMDA and AMPA receptors are involved in the generation of single channel / network bursting and that their balance must be maintained to preserve the network dynamics (Heikkila et al., 2009, Odawara et al., 2016, Frega et al., 2019a, Suresh et al., 2016).

Previous studies regarding the electrical response of dissociated cultured network (Chiappalone et al., 2006, Pan et al., 2009, Eytan et al., 2003, Yvon et al., 2005) shows that two phases of the electrical response can be distinguished. The first phase (i.e. early response) occurs within the first 80 ms and is related to the response of the subpopulation on neurons that are directly excited from the electrical stimulation. The late phase occurs after the 80 ms and was found to be mediated mainly from inhibitory connections that propagate the stimulation the entire network (Chiappalone et al., 2006, Pan et al., 2009, Eytan et al., 2003, Yvon et al., 2005). In our model in which GABAergic neurons are not present, we have found that the response to the stimulus is mainly mediated by AMPA receptors. In fact, AMPA receptors drive the fast dynamics, while on the contrary the NMDA receptors drive the slower dynamics (Rao and Finkbeiner, 2007, Yvon, 2005 #624)]. Under the effect of CNQX the neurospheroids modified the response to the electrical stimulus, not only by decreasing the amount of evocated spikes, but also by changing their response shape. The maximum response peak shifted below 20 ms, because only neurons in close contact with the stimulating electrodes were elicited and they were unable to communicate quickly with others, resulting in failing to mediate the response to the whole network. Following the APV treatment, response to the stimulus changes, finding a lower presence of evoked spikes in the first 100 ms and a delay in reaching the maximum peak. The inhibition of spontaneous fast activity allows for a more precise estimation of propagation speeds and the identification of longer trajectories. This might be cause by the removal of the background noise coming from excitatory postsynaptic potentials, and therefore the average of the raw data centered on a single unit sorted into a channel was less influenced by the activity of other neurons in the surroundings (Bakkum et al., 2008).

Kainic acid is known to cause convulsions in *in-vivo* rats (Sperk, 1994) and it is utilized to model epileptic-like events *in-vitro* (i.e. intense initial bursts followed by repetitive after-discharges) (Vedunova et al., 2013, Colombi et al., 2013, Fisher and Alger, 1984, Odawara et al., 2016, Jimbo and Robinson, 2000, Mzezewa et al., 2022, Odawara et al., 2018). The general effect of KA in the activity exhibited by the neurospheroids was a significant lowering of the number of active electrodes and a consistent increase in the NBR. Furthermore, the response to the stimulus appeared flatter when compared to that in baseline conditions, and the response peak is squashed in the first 20 ms. These two aspects are in line with previous findings on 2D human-derived neuronal network (Odawara et al., 2018, Mzezewa et al., 2022). The connectivity analysis revealed that the electrical stimulation during KA strengthened the connections, by decreasing the number of link, increasing their weight and decreasing their lag. These results suggest that epileptic-like events can be successfully induced in our neurospheroids with KA but further studies are needed to confirm that this is a valid model for studying KA-induced seizures. In fact, all previous works (Vedunova et al., 2013, Colombi et al., 2013, Fisher and Alger, 1984, Odawara et al., 2016, Jimbo and Robinson, 2000, Avoli, 2014) have used KA in cultures where GABAergic neurons were present while we are describing, for the first time in our knowledge, the results deriving from a purely excitatory 3D culture (Mzezewa et al., 2022).

The study of the connectivity *in-vitro* is known to be essential to understand basic mechanism of memory and learning (Poli et al., 2015, Chiappalone et al., 2008, Sporns, 2018, Shahaf and Marom, 2001). The high resolution of HD-MEAs allowed us to evaluate for the first time the connectivity maps of neurospheroids *in vitro*. The inhibition of AMPA receptors increased the disorganization of the network by increasing the number of connections and decreasing their weight. We observed that inhibiting fast transmission greatly decreases the number of connections and only those with heavier weights are preserved. The inhibition of NMDA receptors also led to a decrease in the number of links with an increase in the weight of the connections but in a less clear way. We also recorded a decrease in the average lag of these connections a clear sign that most connections under spontaneous conditions are fast connections mediated mainly by AMPA. The stimulation in the presence of APV instead led to a consolidation of the connections as the number of links decreased, preserving only the connections with the highest weight. This gives a further confirmation of how AMPA and NMDA receptors are also involved in the processes of plasticity and memory (Diering and Huganir, 2018) (Haselmann et al., 2018) (Collingridge, 1987, Li and Tsien, 2009, Rao and Finkbeiner, 2007) and makes our model usable also for this type of study (le Feber et al., 2010, Shahaf and Marom, 2001, Jimbo et al., 1998, Eytan et al., 2003)

Concluding, we have shown how to generate cortical excitatory neurospheroids that can be potentially engineered at will, without the use of scaffolds or bioreactors, and that can be coupled to tools for functional analysis. The fact of generating the neurospheroid starting from differentiated cells instead of pluripotent allowed to generate structures with low variability regard dimension. Since stem cell-derived neurons often require very long maturation times, it is more convenient working with already differentiated cells. In addition development in a 3D conformation induces a morphology more similar to the *in-vivo* in the cells and neurospheroid can be handled and moved at will. The electrophysiological characterization of the spontaneous and modulated / stimulated activity in combination with confocal images indicated that the model presented holds great potential for different applications, ranging from in-depth study of signal transmission to drug screening and disease modeling.

## Supporting information

Supplemental figure with caption

## Author Contributions

Conceptualization, Lorenzo Muzzi; Data curation, Lorenzo Muzzi and Donatella Di Lisa; Formal analysis, Lorenzo Muzzi, Donatella Di Lisa and Matteo Falappa; Investigation, Lorenzo Muzzi and Donatella Di Lisa; Methodology, Lorenzo Muzzi, Donatella Di Lisa and Matteo Falappa; Resources, Sara Pepe, Alessandro Maccione, Laura Pastorino, Monica Frega and Sergio Martinoia; Software, Lorenzo Muzzi; Supervision, Monica Frega and Sergio Martinoia; Validation, Lorenzo Muzzi, Monica Frega and Sergio Martinoia; Visualization, Lorenzo Muzzi, Donatella Di Lisa, Matteo Falappa, Sara Pepe, Alessandro Maccione, Laura Pastorino, Monica Frega and Sergio Martinoia; Writing – original draft, Lorenzo Muzzi, Donatella Di Lisa, Matteo Falappa, Sara Pepe, Alessandro Maccione, Laura Pastorino, Monica Frega and Sergio Martinoia; Writing – review & editing, Lorenzo Muzzi, Donatella Di Lisa, Matteo Falappa, Sara Pepe, Alessandro Maccione, Laura Pastorino, Monica Frega and Sergio Martinoia.

## Notes

Conflict of interest statement: Alessandro Maccione is co-founder and employee of 3Brain AG, Matteo Falappa was a former employee of 3Brain AG.

