## Supplemental figure with caption for "Human-derived cortical neurospheroids coupled to passive, high-density and 3D MEAs: a valid platform for functional tests"

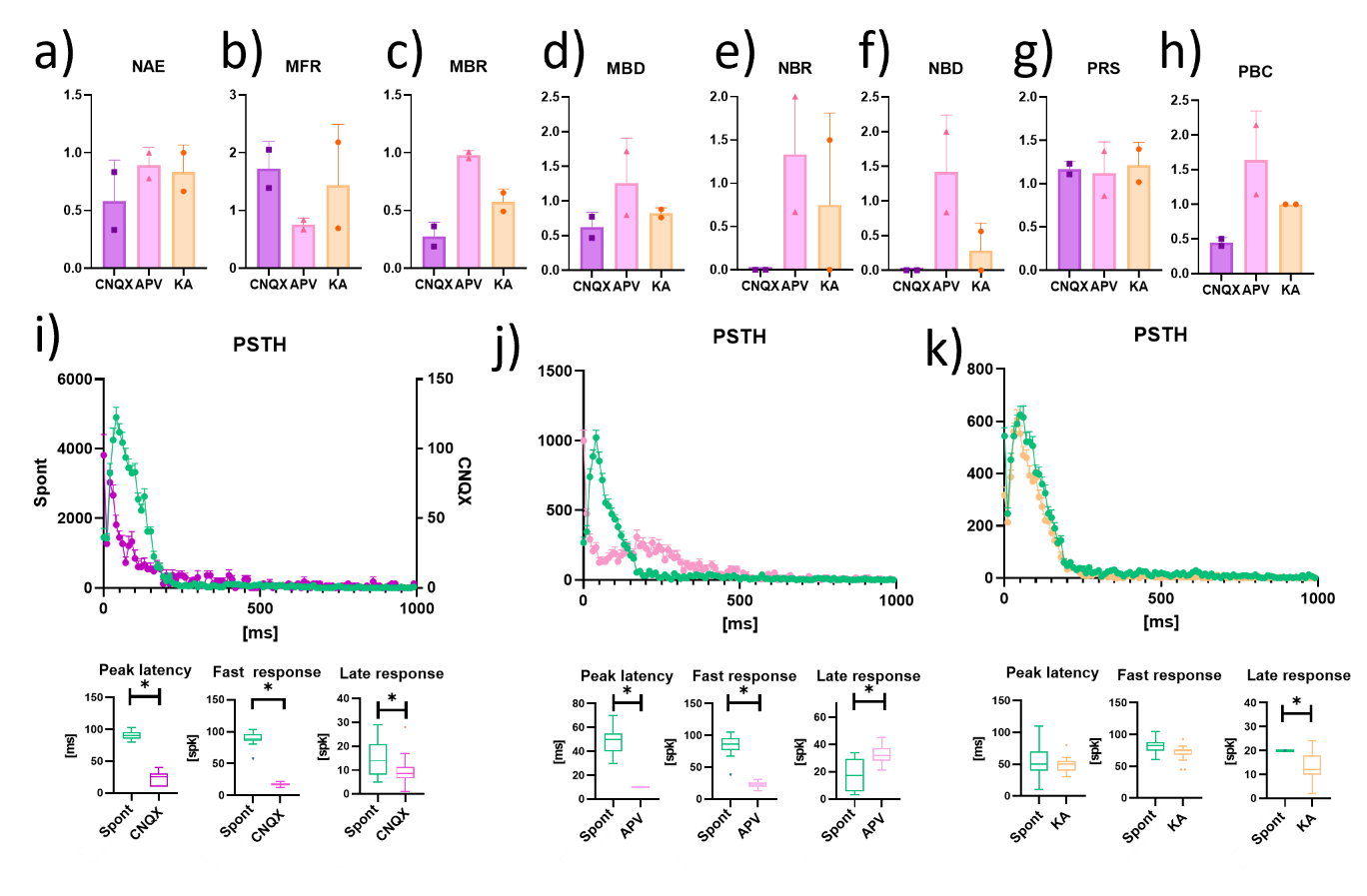

**Suppl. Figure 1.** Evaluation of the electrical and chemical modulation (50 µM CNQX, 80 µM APV, and 5 µM KA). Data are obtained from experiments on MEA60 (n = 2) and are normalized to the value detected in spontaneous conditions in the absence of drugs.. **a-h)** Graphs showing the **a)** Number of active electrodes (NAE), **b)** Mean firing rate (MFR), **c)** Mean bursting Rate (MBR), **d)** Mean burst duration (MBD), **e)** Network Burst rate (NBR), **f)** Network Burst duration (NBD), **g)** Percentual random spikes (PRS), **h)** Percentual bursting channels (PBC) in neurospheroids treated with CNQX, APC and KA, represented int purple, pink and orange, respectively. Data are represented as mean and standard deviation of the mean. **i-k)** Graphs showing the effect of the modulation on the response to the electrical stimulus induced by **i)** CNQX, **j)** APV, **k)** KA. In each panel the Post-Stimulus Time Histogram (PSTH) showing the response of neurospheroids to stimulation is shown (not treated, treated with CNQX, APV and KA are represented in green, purple, pink and orange, respectively). The peak latency represents the mean latency to reach the maximum peak in the PSTHs, the fast response box plot represents the amount of evocated spikes in the first 100 ms of the PSTH while the late response box represent the amount of evocated spikes between 100 ms and 1000 ms. Asterisk indicate p<0.05.
