## Supplementary figures and images for "Human-derived cortical neurospheroids coupled to passive, high-density and 3D MEAs: a valid platform for functional tests"

### Supplemental Figure1

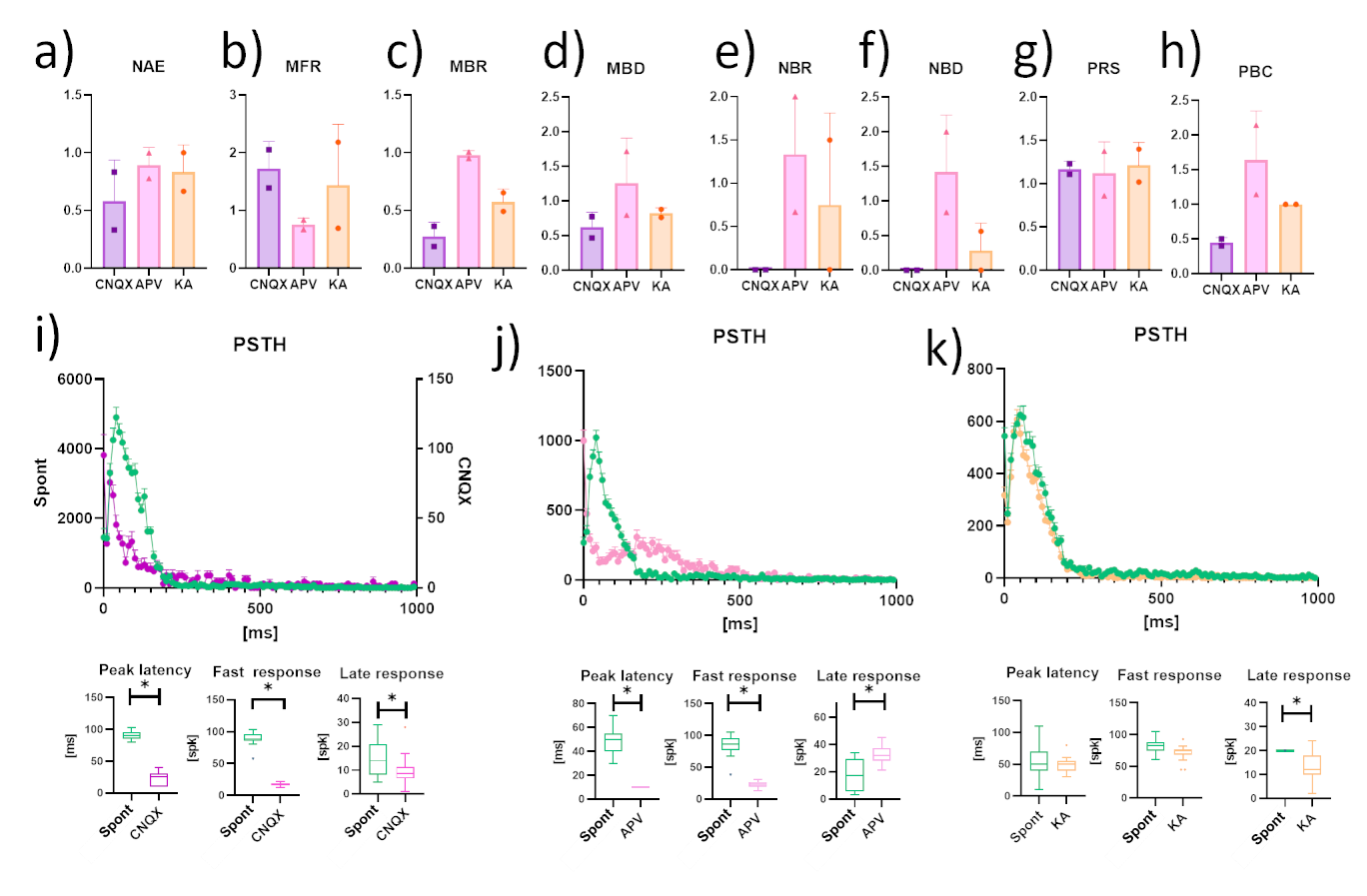
